# Sodium-powered stators of the bacterial flagellar motor can generate torque in the presence of phenamil with mutations near the peptidoglycan-binding region

**DOI:** 10.1101/507533

**Authors:** Tsubasa Ishida, Rie Ito, Jessica Clark, Nicholas J Matzke, Yoshiyuki Sowa, Matthew AB Baker

## Abstract

The bacterial flagellar motor (BFM) powers the rotation that propels swimming bacteria. Rotational torque is generated by harnessing the flow of ions through ion channels known as stators which couple the energy from the ion gradient across the inner membrane to rotation of the rotor. Here we used error-prone PCR to introduce single point mutations into the sodium-powered *Vibrio alginolyticus / Escherichia coli* chimeric stator PotB and selected for motors that exhibited motility in the presence of the sodium-channel inhibitor phenamil. We found single mutations that enable motility under phenamil occurred at two sites: 1) the transmembrane domain of PotB, corresponding to the TM region of the PomB stator from *V. alginolyticus*, and 2) near the peptidoglycan (PG) binding region that corresponds to the C-terminal region of the MotB stator from *E. coli.* Single cell rotation assays confirmed that individual flagellar motors could rotate in up to 100 µM phenamil. Using phylogenetic logistic regression, we found correlation between natural residue variation and ion source at positions corresponding to PotB F22Y, but not at other sites. Our results demonstrate that it is not only the pore region of the stator that moderates motility in the presence of ion-channel blockers.

## Introduction

Motility imparts a large benefit to organisms competing for sparse nutrients. The bacterial flagellum is the oldest known form of motility (Rossmann and Beeby 2018), consisting of a propeller-like filament that rotates under the power of the bacterial flagellar motor (BFM). The BFM is 40 nm in diameter, powered by ion transit across the cell membrane through the stator protein complex, a heterodimer which forms a selective ion channel that transduces chemical energy into mechanical torque (Minamino et al. 2018). While most stators are proton powered (Sowa and Berry 2008), some from marine habitats are sodium powered (Yorimitsu and Homma 2001), others are powered by both sodium ions and protons (Paulick et al. 2015) and recently some have been discovered that are even powered by large divalent cations (Imazawa et al. 2016). In general, flagellar motors have adapted to function in various environments where bacteria live and survive (Terashima et al. 2017; Rossmann and Beeby 2018; Chaban, Coleman, and Beeby 2018). BFM diversity creates an ideal case study to investigate how macromolecular complexes adapt to different environments.

In *E. coli* the energy from the proton gradient is harnessed by the transit of protons through a heterodimeric stator complex of a homotetramer and a homodimer (MotA_4_MotB_2_) (Minamino et al. 2018). In *Vibrio* species this heterodimer is the sodium ion-powered PomA_4_PomB_2_. The total complex, in both cases, consists of four transmembrane (TM) domains of the A-subunit, a single TM domain of the B-subunit, and a large periplasmic region of the B-subunit which consists of a plug segment and a peptidoglycan-binding (PG-binding) region (Kojima and Blair 2001; Kojima et al. 2009). Recently, the structural rearrangements in PG-binding and stator activation were resolved, indicating that conformational rearrangements of the linker between the PG-binding region and the TM-domain are critical in stator activation (Kojima et al. 2018).

Whilst crystal structures have been solved for the C-terminal domains of MotB in multiple species (Roujeinikova 2008; Kojima et al. 2009; Zhu et al. 2014), the full structure of the TM-domain in any species has yet to be resolved beyond low-resolution cryoEM of the entire stator complex (Yonekura, Maki-Yonekura, and Homma 2011). Much of our understanding of stator function in the TM-domain relies on mutagenesis (Sharp, Zhou, and Blair 1995; Toshiharu Yakushi et al. 2006). Chimeric B-subunits have been engineered that combine the C-terminal peptidoglycan binding motif of the MotB subunit with the transmembrane domain of PomB that forms the ion-channel (in complex with two subunits of PomA) (Asai et al. 2003) (Fig. 1). These chimeric PomA_4_PotB_2_ stators utilise sodium motive force to drive flagellar rotation and can bind to the peptidoglycan (PG) layer thus enabling *E. coli* to swim under sodium-motive force. They have enabled the function of the motor to be investigated at low sodium concentration, and thus low energisation (Sowa et al. 2005; Lo et al. 2013).

**Figure 1:**
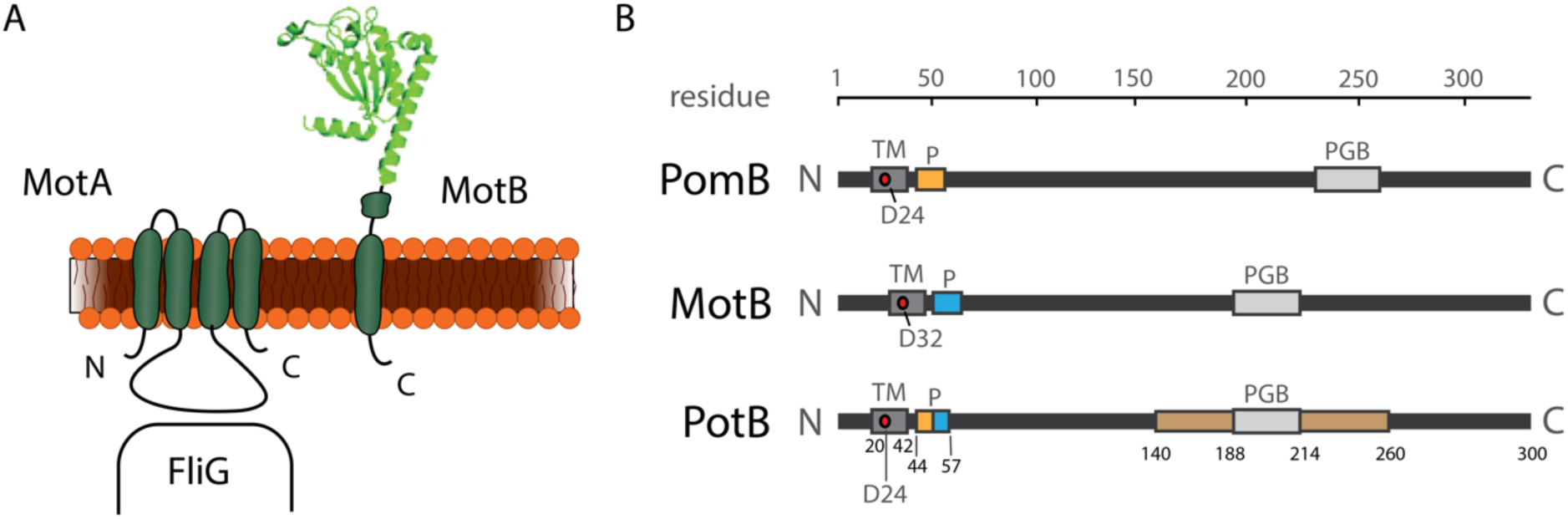
Schematic and structure of the stators of the bacterial flagellar motor. (A) The stator complex in *E. coli* consists of a heterodimeric complex composed of MotA_4_MotB_2_. Each stator has two ion channels that are composed of 4 TM domains of MotA and 1 TM domain of MotB. (B) The PotB chimera consists of residues 1-50 from V. alginolyticus PomA and residues 59-308 of *E. coli* MotB with TM, plug, PGB and ompA-like regions (brown bar) indicated on the schematic respectively. Structure in A from (Seiji Kojima et al. 2008).

Since this chimeric stator complex is driven by sodium ion flow across the inner membrane, it is natural to examine how sodium channel blockers affect this process. Phenamil, a known sodium-channel blocker (Garvin et al. 1985), has been used to probe sodium-interaction sites in sodium driven motors *in situ* in *Vibrio* species (Sugiyama, Cragoe, and Imae 1988; T. Atsumi et al. 1990). Work in *V. alginolyticus* demonstrated that motility under phenamil could be restored by mutations at the cytoplasmic end of the TM-domain of the stator complex (Kojima et al. 1997, 1999). *V. parahaemolyticus* has been widely studied as a model for dual-powered motility; it has lateral flagella that are proton powered but polar flagella that are sodium powered (Tatsuo Atsumi, McCarter, and Imae 1992). Directed evolution approaches have been applied to *V. parahaemolyticus* to examine which spontaneous mutations might induce resistance to phenamil (Jaques, Kim, and McCarter 1999). Jaques et al. selected for resistance to phenamil by selecting flagellar motors that were motile in the presence of 40 µM phenamil methanesulfonate, yet interestingly only observed a single mutant in the TM-domain that resulted in phenamil resistance.

Here we engineered a plasmid construct with enzyme cut sites adjacent to PotB to examine the effects of randomly generated mutations on motility. We screened using phenamil to measure the frequency and location of mutations that enabled motiity under phenamil in the sodium powered PomA_4_PotB_2_ chimera. We induced mutations at a controllable rate using error-prone PCR (error-prone PCR) and screened large populations of cells using streaking of transformed error-prone PCR product onto swim agar in streaks for screening and subsequent sequencing. This allowed a high-throughput screen to determine which mutations have resulted in strains that were functional in the presence of phenamil. We examined how frequently these mutations arose naturally using phylogenetics and parsimony reconstruction to determine whether these adaptations were correlated with existing PomB/MotB classification. By examining the structural and sequence location of these mutations, we have assembled a model that outlines the relationship between PG-binding and stator activity, and presented a framework for determining which residues hold evolutionary importance and may constrain the possible adaptive pathways available to the stators.

## Results

### Mutagenesis of PotB

We generated mutations across the entirety of the PotB protein using error-prone PCR as per Methods. We defined the following regions across the PotB protein (numbering via PotB residue number): the TM domain (20-42), the plug (44-57), the middle domain (57-140) the ompA-like domain (140-260) and the peptidoglycan binding domain (188-214). We screened the combinatoric pool of mutagenesis outputs using swim streaking (Fig. S1) and characterised the motility of 27 error-prone PCR plasmids that exhibited some motility when selected for motility in the presence of phenamil (Fig. S2; Table S1). The distribution of all nucleotide mutations in all 27 plasmids, including silent mutations, is shown in Fig. 2.

**Figure 2:**
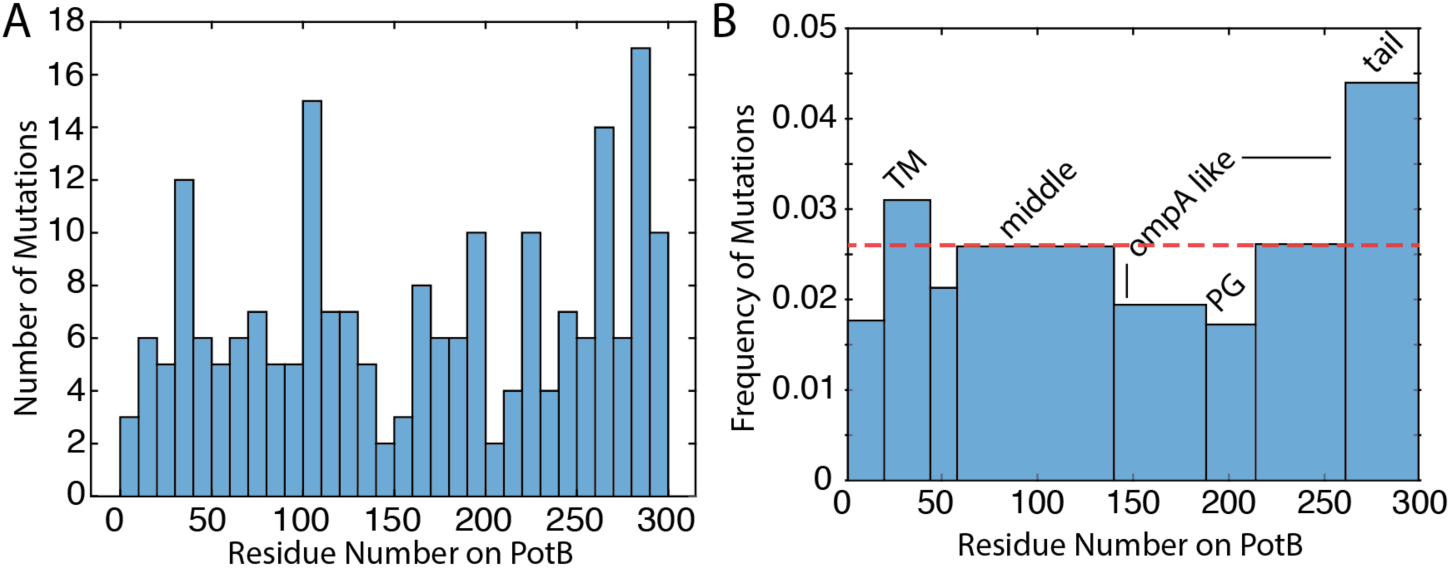
Histogram of mutant frequency. Histograms of mutations from error-prone PCR across every strain with at least partial motility in phenamil (from spread plate). (A) Binned evenly across PotB with each bin 10 residues in width showing raw counts of mutations across aggregated results from all 27 error-prone PCR plasmids. (B) Frequency of mutations binned according to specific regions of PotB: TM (20-42), plug (44-57), middle (58-139), ompA-like first section (140-187), PG region (188-214), ompA-like second section (215-260) and tail (261-300). Counts are normalised across each region and strain (average mutations per residue per strain), and the mean across the entire protein is indicated by the dashed red line (0.026 mutations per residue per strain)

Together, these histograms indicated that the mutations occurred roughly evenly throughout the protein. We quantified the mutation rate in each region of the protein by calculating the mean number of mutations per strain in each region and normalising for the length of each region to calculate a mutation rate per residue for each region (Fig. 2B). We observed that proportionately more mutations occurred in the TM and the tail region of PotB, that is, the mutation rate was 0.031 and 0.044 mutations/residue in the TM and tail region respectively, which was greater than the overall mutation rate across the entire protein, which was 0.026 mutations/residue.

### Point mutations in TM domain can enable motility in the presence of phenamil

Out of the 27 strains (Table S1), 209 mutations were observed in total, with a median number of mutations per strain of 8. Of the 209 nucleotide mutations, 62 were silent mutations that did not result in an amino acid change. Six exhibited a strong motility phenotype when motility was characterised in the presence of 20 µM phenamil. Of particular interest, we focused on plasmid PotB-ep18 (Fig. S1-S2, Table S1) which contained only 3 mutations: F22Y, L28Q, K100Q and demonstrated increased motility in the presence of phenamil. To address the relative contributions of each mutation to the observed phenotype we synthesised single point mutants of each of the three mutations (Fig. S3). Both F22Y and L28Q alone enabled motility under phenamil to PotB (Fig. 3), or together (pSHU149, F22Y/L28Q), where they were slightly additive. Mutation K100Q appeared to offer no benefit in motility under phenamil and as such was classified a redundant mutation. These results indicate that single point mutations, particularly in the pore region of the TM domain of PotB, can enable motility under phenamil.

**Figure 3:**
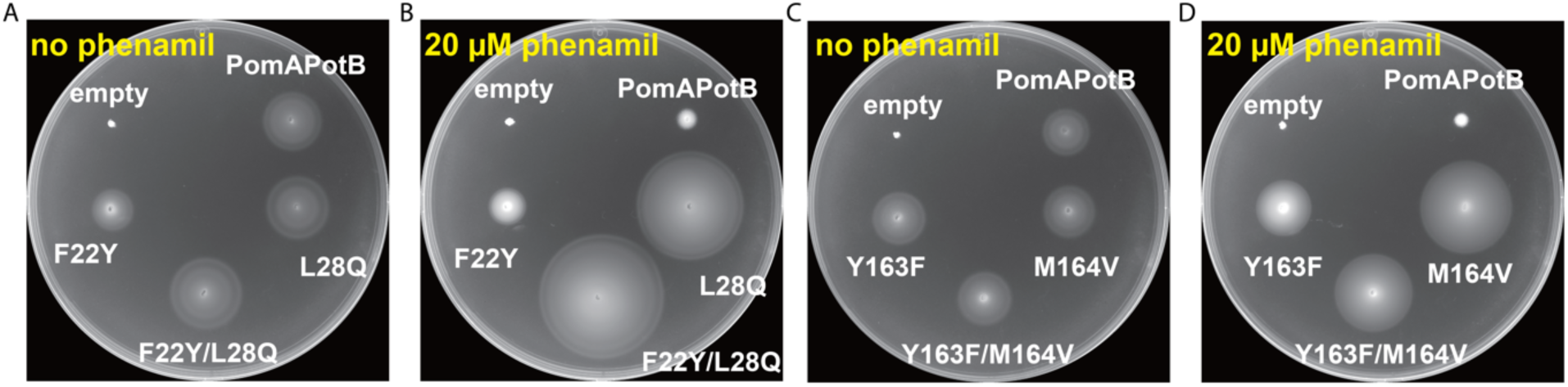
Mutants in TM-domain and PG-binding region enable motility under phenamil. Motility of either TM-domain mutants (A/B) or PG-binding mutants (C/D) was tested via 0.25% agar swim plating using the empty vector (pBAD33) and PomAPotB wild-type as a negative and positive control respectively. A/C) In the absence of phenamil, swimming of mutants is comparable to the positive control. B/D) In the presence of 20 µM phenamil, the mutants are motile whilst PomAPotB wild-type is severely limited. A/C incubated at 30°C for 10 hours, B/D incubated at 30°C for 18 hours.

### Point mutations near PG-binding domain enable motility in the presence of phenamil

Plasmid PotB-ep9 (Fig. S1-S2; Table S1) displayed a phenamil-resistant phenotype and also had only three mutations: Y163F, M164V and L255M. This was targeted for further investigation as these mutations were all far from the TM domain. We again characterised the contributions of each mutation to motility and observed that Y163F and M164V each separately enabled motility in the presence of phenamil and that L255M was a redundant mutation (Fig. S3). When both mutations were combined in a single plasmid (pSHU146), motility in the presence of phenamil was increased (Fig. 3).

### PG-binding double mutants rotate at lower sodium concentration than wild-type PotB

To establish whether these mutants were generating torque from sodium motive force, or had undergone a change in ion-specificity, we tested swimming in minimal sodium-free media (Fig. S4). None of the mutants were motile in sodium-free media, yet all plasmids restored swimming in the presence of sodium and in the presence and absence of phenamil (Fig S4). We then tested individual cells for rotation whilst lowering the concentration of sodium to determine whether the dependence on sodium-motive force (SMF) had changed in these mutants (Fig. 4). This allowed us to confirm that our observed swim plate phenotype was caused changes in flagellar rotation, and account for any compensatory changes in growth or chemotaxis that might confound swim plate measurements. We examined the capacity of ‘wild-type’ PomAPotB (expressed via our plasmid pSHU1234, Fig. S5) and our mutant plasmids to restore swimming in ‘sticky-filament’ strains. These sticky-filament mutants lack the surface of the protein FliC and thus undergo hydrophobic interactions with glass that enable filaments to stick to glass and allow the rotation of the body to be assessed in parallel (Kuwajima 1988). In comparison with PomAPotB wild-type, the mutants in the PG-region (pSHU146: Y163F/M164V) were highly motile at low sodium (further detail Fig. S6), with cells rotating > 6 Hz in the presence of only 0.3 mM Na^+^. This is in contrast to PomAPotB wild-type restoring motility to < 2 Hz at the equivalent sodium motive force. In an opposite trend, mutations in the TM region (pSHU149: F22Y/L28Q) did not demonstrate swimming below 1 mM Na^+^ and had a lower maximum speed and slower response to increased [Na^+^].

**Figure 4:**
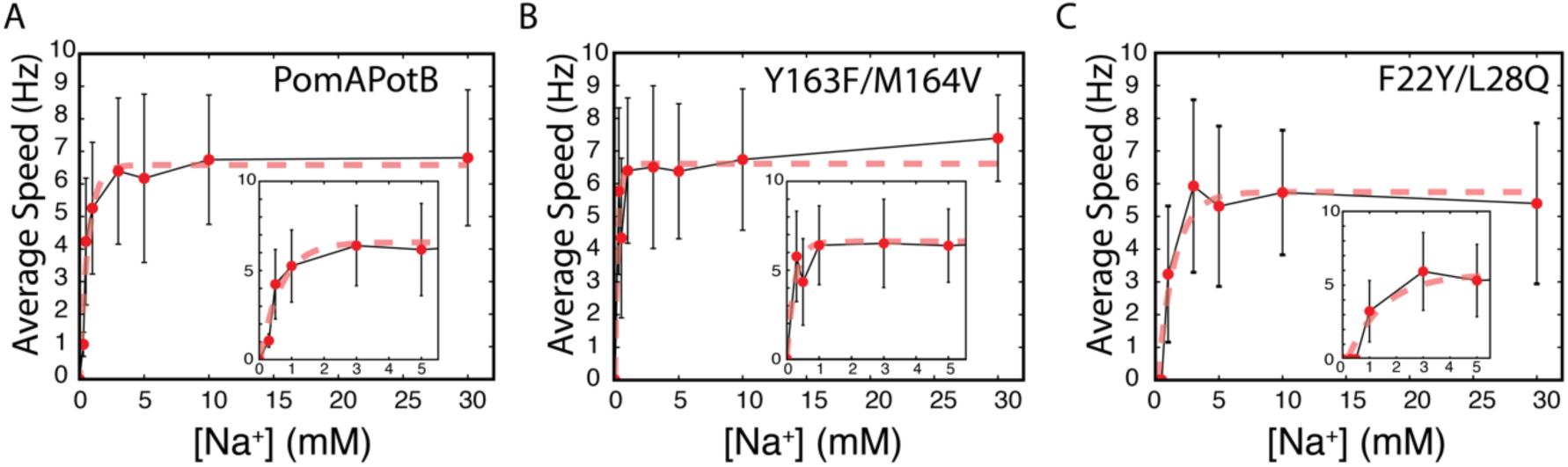
Y163F/M164V is motile at less than 0.3 mM [NaCl]. (A) The PomAPotB wild-type exhibits single cell rotation from 0.3 mM Na^+^, increasing exponentially to a plateau above 6 Hz at full energisation. (B) PotB Y163F/M164V displays rotation at 0.3 mM Na^+^, with plateau above 6 Hz at full energisation. (C) potB F22Y/L28Q displays no rotation below 1 mM Na^+^, with a slower increase to a lower maximum speed plateau above 5 Hz and under 6 Hz. Inset in ABC shows zoom around 0-5 mM [Na^+^]. Dashed lines indicate monoexponential fit (ω = *ae*^-*bx*^ + *c*).

**Figure 5:**
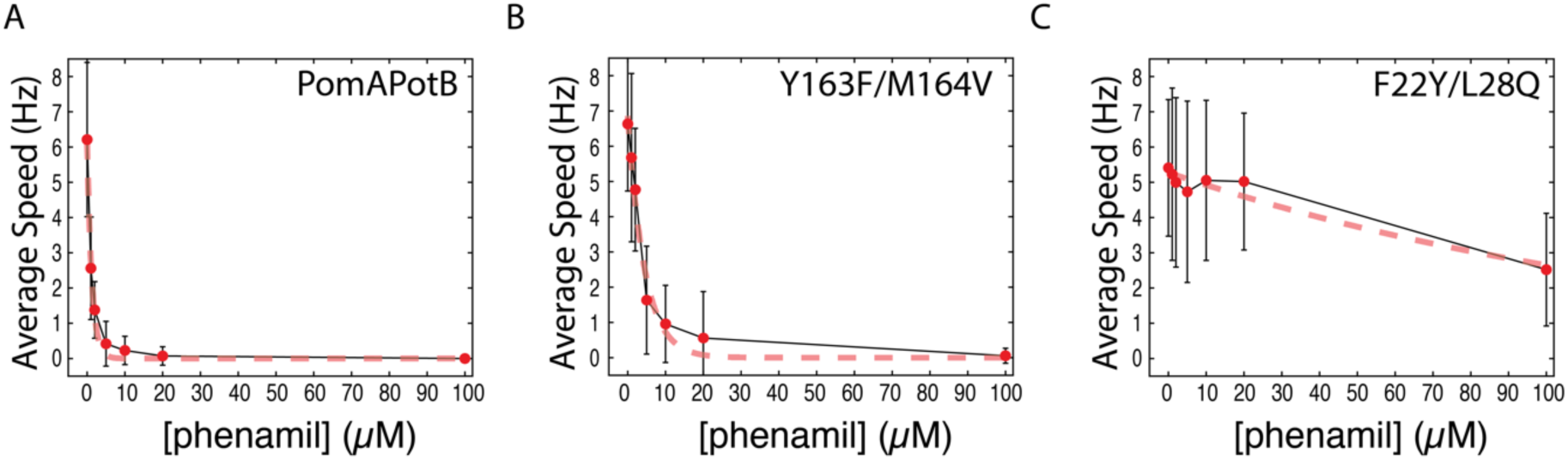
TM mutants enable motility in phenamil concentrations up to 100 µM. Dose response of averaged measurements of single-cell rotation speed with varying concentration of phenamil for: (A) PomAPotB wild-type, double TM-domain mutant (PomAPotB F22Y/L28Q) and (C) double PG-binding mutant (PomAPotB Y163F/M164V). Tethered cell data is fit with monoexponential decay curve to generate time constants for decay (PomAPotB wild-type: 0.8 mM^-1^; PomAPotB Y163F/M164V: 0.2 mM^-1^; PomAPotB F22Y/L28Q: 0.007 mM^-1^).

### TM-domain double mutants are not affected by increasing concentrations of phenamil

The response of single-cell speed with varying phenamil concentration was measured to examine the energy profile and structural basis of motility under phenamil in our two phenamil-resistant plasmids. In wild-type PomAPotB tethered cell rotation speed was reduced significantly when as little as 1 µM of phenamil was introduced, and cell speed plateaued at 5 µM of phenamil. The PG-binding double mutant (Y163F/M164V) demonstrated the same plateau effect in response to increasing phenamil but the concentration dependence of the decrease in rotational speed was much lower, that is, the decay constants for a monoexponential fit (fitting for *b* in: ω = *ae*^-*bx*^) were 0.8 mM^-1^ and 0.2 mM^-1^ for PomAPotB wild-type and Y163F/M164V respectively. Similarly, the I_50_, the concentrations of phenamil at which the speed was half the maximum, was 0.9 µM and 3.1 µM for wild-type and Y163F/M164V respectively. In contrast, the speed of rotation for the double TM-domain mutant (F22Y/L28Q) exhibited no dependence on phenamil concentration up to 20 µM, and only a 46% reduction on maximum speed at 100 µM phenamil. This implies that the mechanism of resistance in the TM-domain mutants destroyed much of the efficacy of phenamil to inhibit motility altogether, whereas the PG-binding region mutants acted to stabilise rotation in the presence of phenamil, but this stabilisation could be overcome as phenamil concentration was increased.

### F22Y is common and clusters in phylogenies along MotB/PomB lines; L28Q is very rare

To explore the natural frequency of these residues and their correlation with ion specificity and motility outside the lab, a large low-level phylogeny of 948 MotB homologues was assembled which clearly divided into clades (Fig. 6A). We curated a subset of 34 commonly studied species where information on ion specificity exists in the literature (Fig. 6BCD). For each column of the subset alignment, we reconstructed ancestral residues with parsimony. At the F22Y site, residues clustered according to MotB/PomB identity (Fig. 6B), which is putatively linked to proton/sodium motility. To test if the correlation is greater than expected by chance, we used generalized linear models to perform phylogenetic and non-phylogenetic logistic regression (see Methods). This demonstrated that F/Y at position 22 had significant correlation with PomB/MotB respectively (Supplementary Table S3). In contrast, at L28Q (Fig 6D), the residue variation had no significant correlation with PomB/MotB. In fact, over the entire 948 species in the larger phylogeny, only one species had a glutamine at that site: *Myxococcus fulvus.*

**Figure 6:**
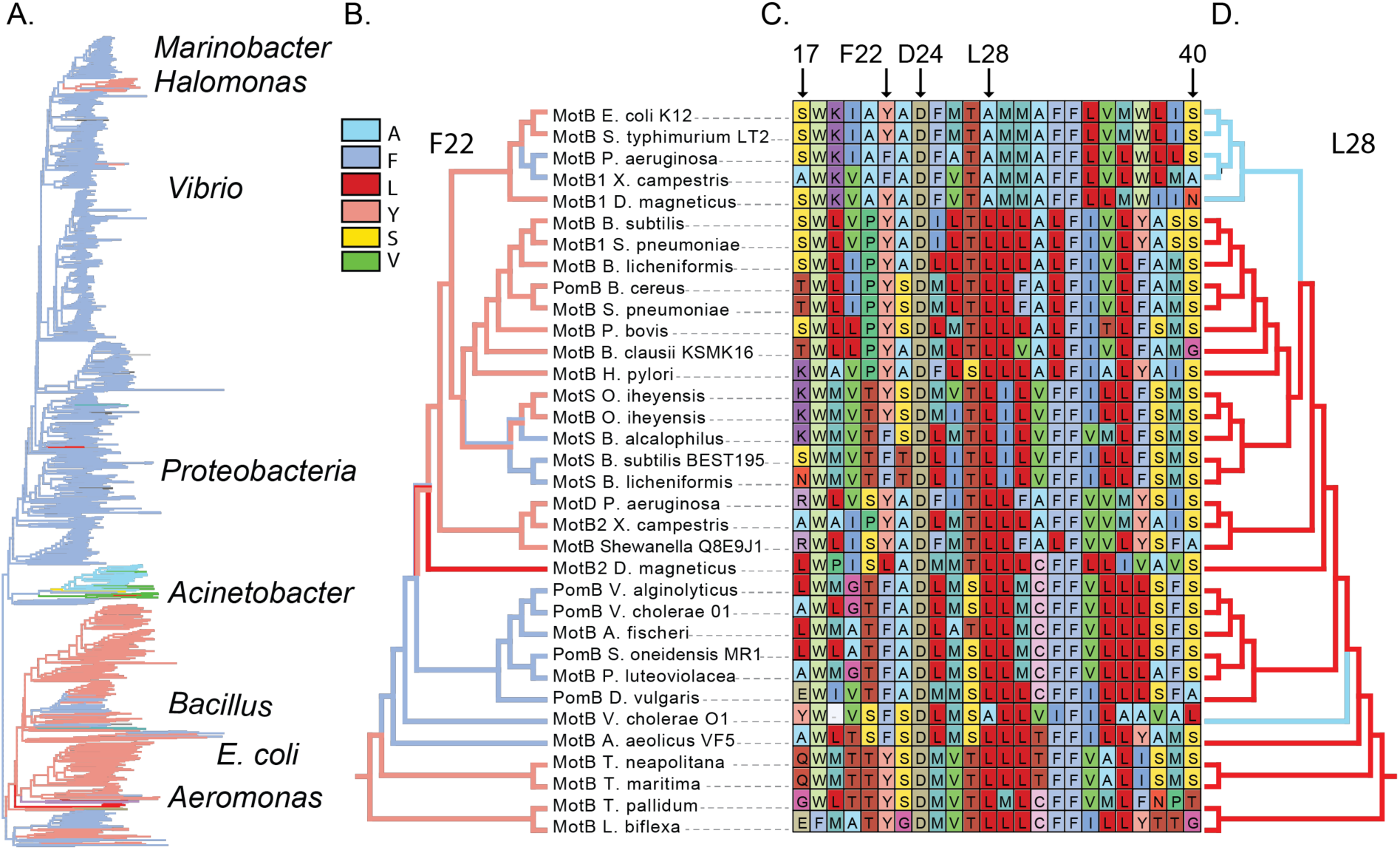
Parsimony Reconstruction for Y22 and L28. (A) Overall phylogeny of 948 MotB homologs coloured via sequence identity at site 22 showing clear signal (well-known clades are reproduced). Some major phyla indicated. (B) Parsimony reconstruction for site Y22 (PotB numbering) shown in colour over subset phylogeny of 34 representative species as labelled. (C) Sequence alignment across transmembrane domain of MotB/PomB showing conserved D24, and Y22/L28 mutations tested in this work. (D) Parsimony reconstruction for site L28A (PotB numbering) shown in colour over subset phylogeny.

### Y163F/M164V are both rare and do not cluster along MotB/PomB lines

Similarly, we aligned the sequences in the section 156-214, including the PG-binding site, and measured correlation with PomB/MotB (Fig. 7, Table S3). In both Y163F and M164V, logistic regression could not be run as these columns are not predominantly composed of 2 residues (our algorithm requires a binary predictor variable). However, visual inspection suggests the mutations do not cluster along PomB/MotB lines. A phenylalanine at site 163 was very rare across the full phylogeny, occurring only in *Yersinia pestis biovar Orientalis.* The valine at site 163 was observed in 13 species, mostly among members of the *Spirochaetes* and *Oceanospirilla* families (Supplementary Note).

**Figure 7:**
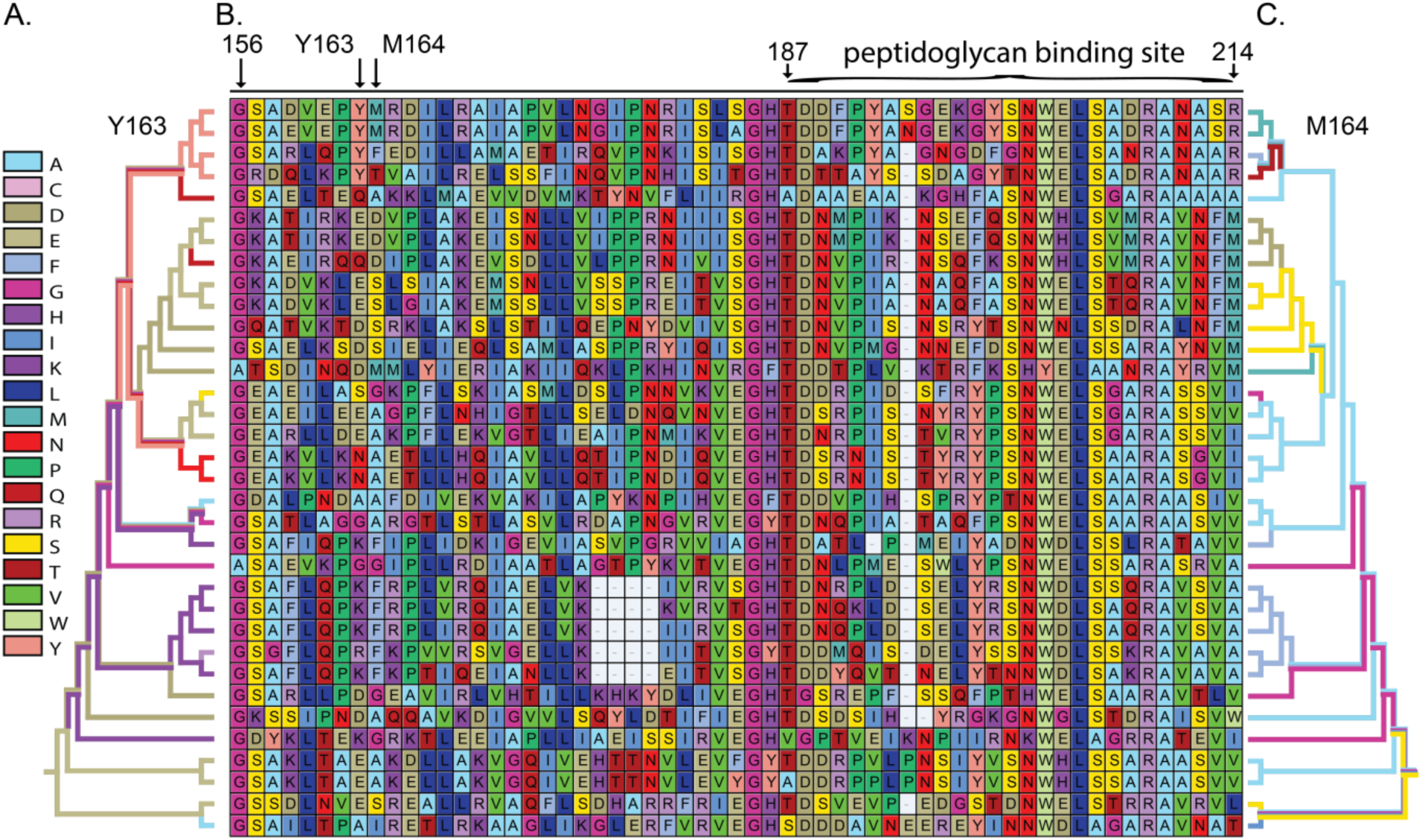
Parsimony Reconstruction for Y163 and M164. (A) Parsimony reconstruction for site Y163 (PotB numbering) shown in colour over subset phylogeny of 34 representative species from Fig. 5. (B) Sequence alignment over peptidoglycan binding site of MotB/PomB showing Y163/M164 mutations tested in this work. (C) Parsimony reconstruction for site M164 (PotB numbering) shown in colour over subset phylogeny.

## Discussion

We have examined the effect of mutations in both the transmembrane domain and the peptidoglycan binding region of PotB to the binding of phenamil. Phenamil binding has been suggested to bind on the cytoplasmic face of the stator(S. Kojima et al. 1997, 1999; T. Yakushi, Kojima, and Homma 2004), and function by limiting Na^+^ exit into the cell interior (Yoshida et al. 1990). However, the question remains how the mutations observed here, far from the pore, might influence phenamil binding and sodium transit. We observed that single point mutations in the TM region at L28 and F22 were sufficient to enable motility under phenamil. The mutation at F22Y is highly clustered around Pom/Mot stator subunits in varying species and presumably linked to sodium/proton specificity, which may show that residue F22 not only plays a role in sodium binding but also phenamil-binding and sodium blocking. Together with a mutation we observed as an error-prone PCR product at L28Q, these two mutations here have enabled motility under phenamil across a large range of phenamil concentrations, with stators remaining functional at concentrations of phenamil as high as 100 µM phenamil. This high tolerance to phenamil implies that they have affected or removed the binding site of phenamil inside the pore region of the stator complex.

Previous work in *V. Alginolyticus* extensively explored the effect of mutations at PomB site 22 on motility in the presence and absence of phenamil (Terauchi et al. 2011). Of these point mutations in PomB (as opposed to chimeric PotB), the substitutions F22S, F22W and F22N provided high motility under phenamil, however it was also observed that F22Y displayed a phenotype of motility under phenamil. Our results confirm the importance of F22 site in the chimeric PotB and likely proximity to the Na^+^ binding site and involvement in the ion-conducting pathway, and show that even slight changes in the size of this residue can affect the binding site and enable motility under phenamil.

Similarly, in the PG-binding region we observed that two clustered single mutations were also sufficient to enable motility under phenamil. The PG-binding region is used to bind the stators in place so that they act on the motor. With these two mutations at Y163/M164 we observed a decaying concentration dependence of rotation to phenamil. These sites are far from the pore, which is expected to be the phenamil binding site. We consider that mutations near the PG region could allow motility under phenamil via two models: 1) changes to the PG region can impact stator-rotor interface and alter the pore region and phenamil binding site; and 2) that changes to the PG region can alter the binding strength of PotB to the PG layer in *E. coli* and alter the dwell time of the stator on the rotor. The first option is supported by previous evidence that a mutation in the PG region of MotB (P159I) could be suppressed by a mutation on the rotor protein FliG (K192E) (Garza et al. 1995). This suggested that changes in the PG region could influence the rotor-stator interface, and that this interface could be realigned with a compensatory mutation in the rotor. Thus it is conceivable that our observed mutations in the PG region (Y163/M164) are influencing the rotor-stator interface and thus altering the ion channel and the phenamil binding site, in a similar mechanism to changed directly at the binding site caused by L28Q and F22Y. The second model contends that a change in the binding strength of the stator might increase the dwell time of the stator on the rotor is result in more stators being bound more often (Fig. 8). In turn, if each stator is fractionally functional, or occasionally able to pass a sodium ion in the presence of phenamil, then in total, with more stators on the rotor, these sodium-powered flagellar motors are able to function in the presence of phenamil. Recent structural evidence indicates that upon binding to the PG layer there is a dynamic rearrangement where a helix (helix *α*1) is extended into a non-helical structure to activates the stator and open the ion channel (Seiji Kojima et al. 2018). Thus it is also conceivable that mutations in the PG region could affect the release from the PG layer, the extension of helix *α*1, or the conformational change that removes the plug from the channel. Further experiments are needed to distinguish between these models, for example, the use of single cell studies and single molecule fluorescence to measure changes in the dwell time of mutant stators on the motor, and tight regulation of stator expression to measure for dependence of motility on stator type, concentration and availability.

**Figure 8:**
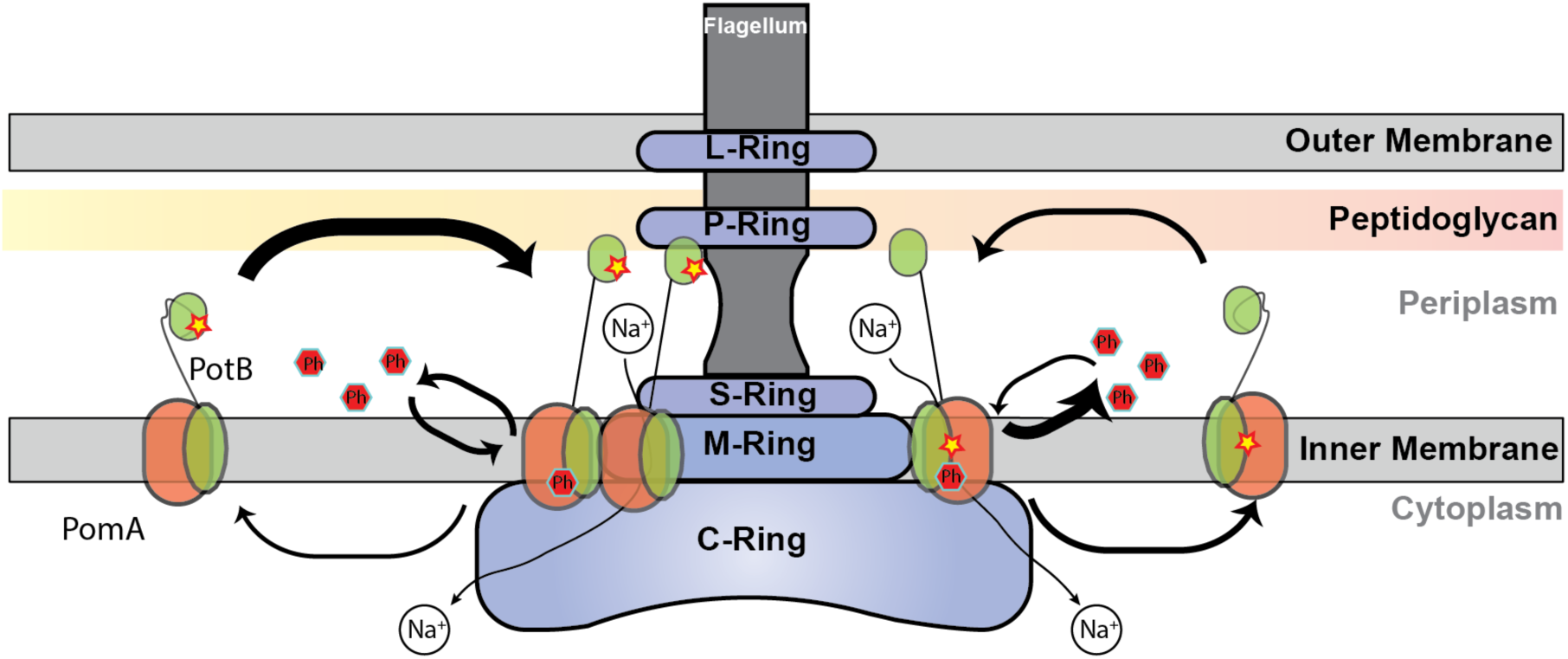
Model of activity. Schematic showing effects of mutations with respect to phenamil and sodium flow through stator complex. The pool of stator complexes (PomA_4_PotB_2_) diffuse in the inner membrane, and when they bind to the motor the PG-binding region pulls with tension on the transmembrane domains of PomA and PotB (Kojima et al. 2018). (left) Mutations in the 163/164 site affect PG-binding and the duration for which a stator is bound to the motor (dwell time). This could result in a slower off rate with respect to wild-type stators and a larger steady-state population of stators bound to the motor. In the presence of phenamil each stator has reduced functionality but in sum, with many stators, the motor is functional. (right) Mutations at the TM-domain near the putative phenamil binding site affect the binding and release of phenamil, thus phenamil is bound more weakly, and turns over more rapidly, allowing sodium ions to pass through the channel to generate motor rotation. An alternate possibility is that that mutations at 163/164 site cause a consequential alteration of the binding site and thus limit phenamil binding in a similar manner to mutations at 22/28 site.

Phenamil resistance has been studied in other species, most notably in *Vibrio parahaemolyticus*. This system is of interest as it has two motors, one sodium-powered and one proton-powered and phenamil has been studied previously as a drug target for this pathogen (Tatsuo Atsumi, McCartert, and Imae 1992; Jaques, Kim, and McCarter 1999). Jaques et al. used natural selection over 12-18 days to screen for phenamil resistance (Jaques, Kim, and McCarter 1999). They observed a single mutation in the TM-domain, but did not observe the mutations that we saw in this paper. Their experiment, occurring through natural mutagenesis and evolution in a laboratory setting will respond to pleiotropy and epistasis and also inherent directionality in the process of stator adaptation. This means that the order of specific mutations can restrict the possible outcomes. Naturally, our mutation at L28Q was indeed very rare across our set of 948 MotB homologues, occurring only once in *Myxococcus fulvus* ∽0.1% of the species we examined. Error-prone PCR however, is agnostic to the order of mutations and will insert mutations evenly across the protein of interest. This highlights the power of error-prone PCR to explore novel mutations and to compare historical evolutionary occurrences with those that can be discovered via directed and laboratory-assisted experimental evolution.

Our experimental system allows us to examine phenotypes to determine which amino acids have a functional cause in changing a phenotype. This allows us to address a common puzzle in phylogenetics: because both mutations and traits are inherited on the phylogeny, they are non-independent, and can thus exhibit “accidental” correlation without a functional relationship (Uyeda, Zenil-Ferguson, and Pennell 2018). For example, in the case of MotB, every clade has some amino acids unique to it, but this will be true whether or not those residues have a functional role in the clade’s motility phenotype. However, by characterising upmotile mutations in the presence of phenamil and adaptations in ion-source, we can directly test whether amino acid substitutions observed on the phylogeny are causally associated with a particular ion source, or a particular environment. This allows us to identify which amino acids, potentially in which order, have been part of historic adaptation events such as a transition from sodium to proton-based motility. We can then use these approaches to tease out the causality of events that constrain natural adaptation.

### Experimental Procedures

#### Bacterial strains and plasmids

Bacterial strains and plasmids used in this study are listed in SI. For swim plating, plasmids were transformed into RP6665 (Δ*motAmotB*) (Block, Blair, and Berg 1989) to restore motility. For tethered cell assays, plasmids were transformed into JHC36 (Δ*cheY fliC-sticky* Δ*pilA* Δ*motAmotB*) (Inoue et al. 2008).

#### Error-prone PCR

Plasmid pSHU1234 was constructed in a pBAD33 backbone with unique cut-sites for NdeI and PstI directly upstream and downstream of PotB. Error-prone PCR was executed on pSHU1234 using primer 1176 and primer 0104 (Table S2), taq polymerase, unbalanced dNTPs and Mn^2+^, as per (Wilson and Keefe 2000). The post-PCR product and pSHU1234 was then digested and ligated with NdeI and PstI and transformed into RP6665(Δ*motAB*).

#### Mutant isolation

Tranformant was streaked in lines on low agarose motility plate (Fig. S1) consisting of 0.25% TB soft agar (0.25% bactoagar, 1% bactotryptone, 0.5% NaCl), 25 µg/mL chloramphenicol, 1 mM arabinose and 20 µM phenamil. These plates were incubated at 30°C for 20 hours. Flares were selected from these plates and single colonies were generated. These colonies were harvested, the plasmid was purified, transformed again into RP6665 and motility was tested as below. Specific mutants derived from error-prone PCR outcomes were engineered using a standard site-directed mutagenesis protocol with forward and reverse primers (Supplementary Materials), template, and high-fidelity DNA polymerase (Pfu Ultra).

#### Motility assays

Motility testing was carried out using motility plate assays. Phenamil to a final concentration of either 0, 20 or 100 µM was added to the plates. Swim plates were inoculated from single colonies by toothpick. Plates were incubated for 10 or 18 hours respectively, as noted. For standard motility assays, plates consisted of minimal media for minimal motility assays (Fig. S4) consisted of 10 mM KPO_4_, 0.1% glycerol, 0.1 mM Thr, 0.1 mM Leu, 0.1 mM His, 0.1 mM Met, 0.1 mM Ser, 1 mM MgSO_4_, 1 mM (NH_2_)_2_SO_4_, 1 μg/mL Thiamine, 0.5% NaCl or KCl respectively (85 mM [NaCl]; [67 mM KCl]), 1 mM arabinose, 25 µg/mL chloramphenicol and the indicated concentration of phenamil (0, 20, 100 µM).

#### Sequence analysis

Sequencing was executed commercially by FASMAC (Japan) using forward and reverse primers and template (Table S2). Three primers were used to read the entire PomAPotB region.

#### Single cell speed assays

Cells were adhered spontaneously to glass coverslips via a sticky-filament and then imaged onto a CMOS camera at 60 frames per second thorough 40x objective. Rotation of tethered sticky-filament cells was analysed using custom software based upon LabView (National Instruments).

#### Phylogenetics and sequence diversity of MotB homologs

To survey the known flagellar diversity for amino acid substitutions similar to the upmotile mutations discovered by experiment, we estimated a very large phylogeny of MotB and related proteins. As our goal was exploratory, we chose faster heuristic methods feasible for a large dataset, rather than Maximum Likelihood or Bayesian methods that would be impracticably slow. 948 MotB homologs were assembled by searching the UniProt90 database against key study taxa. They were aligned using Clustal Omega (Larkin et al. 2007) on its most thorough settings (5 iterations of re-alignment). The phylogeny was then estimated with Quicktree, a neighbour-joining method suitable for large datasets (Howe, Bateman, and Durbin 2002). The phylogeny was midpoint-rooted in FigTree (Rambaut 2018). As these are fast heuristic methods based on a single protein, deeper branches and rooting should be considered uncertain, and we make no claim about e.g. inter-phylum relationships, a very difficult phylogenetic problem (Pallen and Matzke 2006; Abby and Rocha 2012; Shih and Matzke 2013; Koonin 2016). However, the tree was adequate for the purpose of showing relationships between closer relatives, and for surveying MotB sequence conservation and diversity.

To further explore conservation and diversity at key positions of the alignment, ancestral amino acids were reconstructed on the phylogeny using parsimony in Mesquite (Maddison and Maddison 2015). The 948-protein tree was subset to key study taxa for Fig. 6 and Fig. 7.

#### Logistic Regression

We classified each species in the 34-species subset phylogeny as Na+, H+, or N/A to create a binary response variable for a logistic regression analysis. We were interested in cases where this binary response might be correlated with a binary amino acid predictor, so we filtered the columns of the subset alignment for columns where only 2 amino acids were predominantly observed. We excluded any column where the two most common amino acids added up to <85% of the observed residues. We also excluded any site where the first or second most common amino acid had a <10% frequency. These steps excluded the 163/164 site, but included positions 22/28. We ran two models to correlate each amino acid or the mutational pair with the ion source available: (1) non-phylogenetic logistic GLM (glm, R), and (2) a phylogenetic logistic glm (phyloglm, R; Ives & Garland, 2010). We present the slope estimates, errors, and *z* scores and *p*-values in Table S4, and our classification list of ionic energy sources in Table S5.

## Author Contributions

TI collected and analysed data. RI collected and analysed data. JC collected data. NJM analysed data and wrote the manuscript. YS conceived and supervised the project, collected and analysed data, and wrote the manuscript. MABB conceived and supervised the project, collected and analysed data, and wrote the manuscript.

