## Supplementary Material for "Sodium-powered stators of the bacterial flagellar motor can generate torque in the presence of phenamil with mutations near the peptidoglycan-binding region"

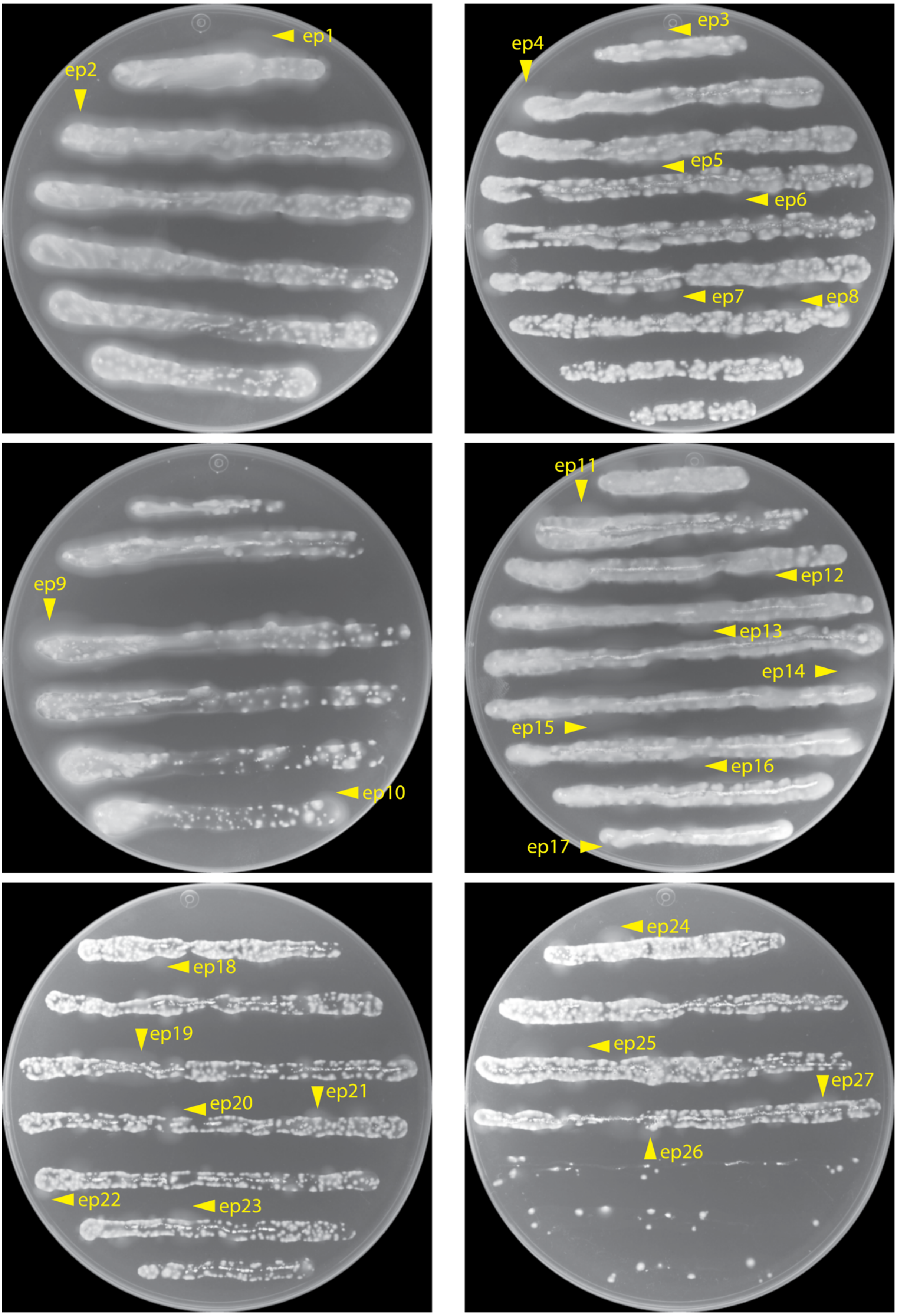

**Figure S1: Streak Plates of EP-PCR Products.** Streak plates used to select the 27 strains identified in this work. Host: RP6665, 0.25% TB agar, 1 mM arabinose, 20 µM phenamil, incubated at 30˚C for 20 hours.

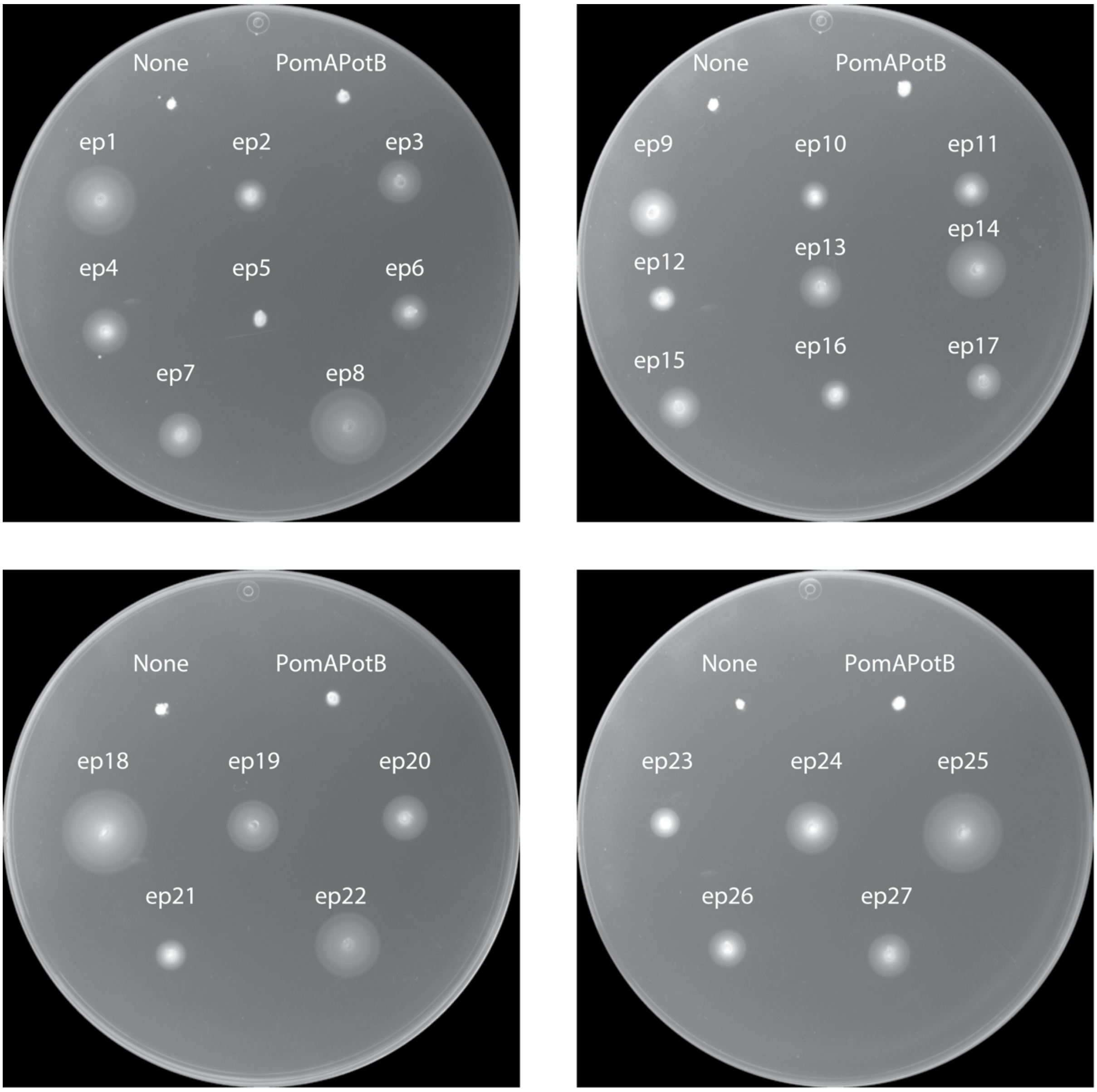

**Figure S2. Motility characterisation of screened outcomes from streak plates.** Motility characterisation for strains identified in streak plates shown in Fig. S1. Host: RP6665, 0.25% TB agar, 1 mM arabinose, 20 µM phenamil, incubated at 30˚C for 11 hours.

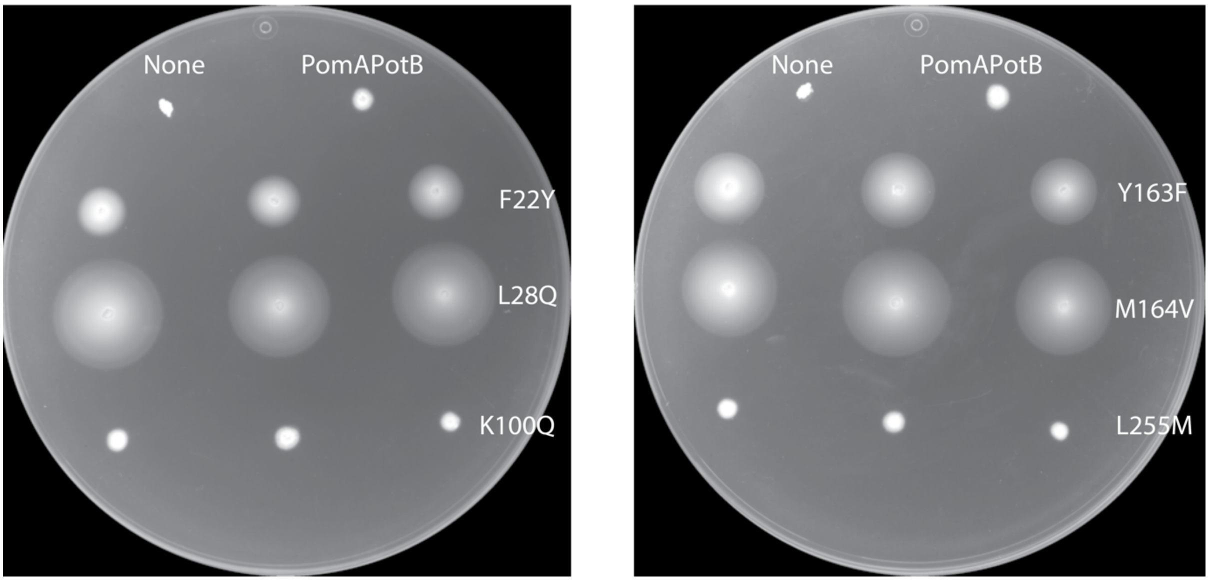

**Figure S3. Characterisation of functional and non-functional mutants in PotB-ep18 and PotB-ep9.** Test of individual contribution of the three mutations in PotB-ep18 (left) and PotB-ep9 (right) respectively. Host: RP6665, 0.25% TB agar, 1 mM arabinose, 20 µM phenamil, incubated at 30˚C for 17.5 hours.

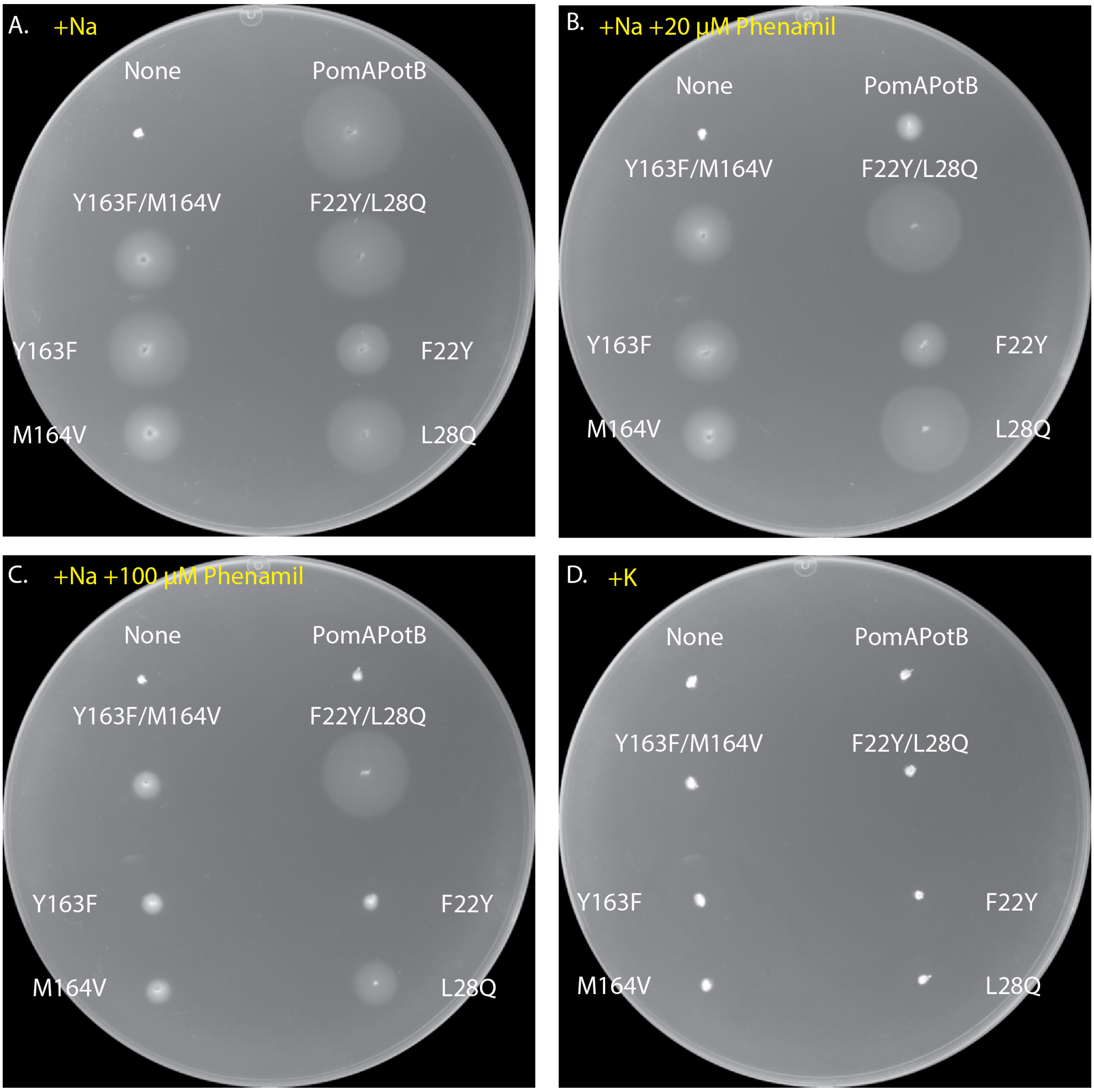

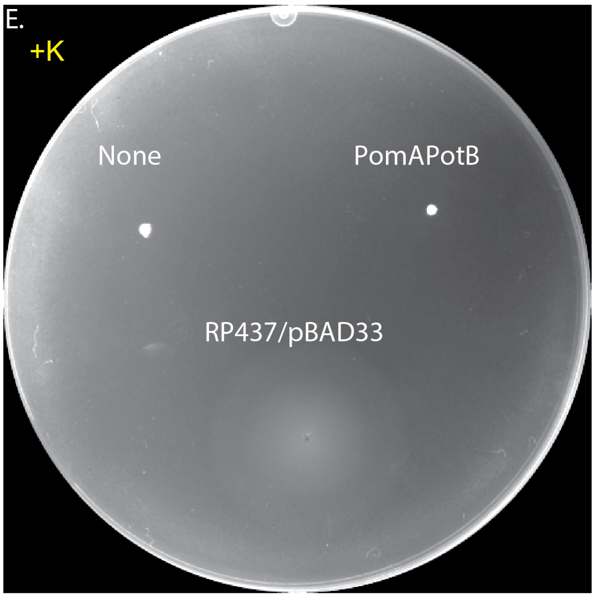

Figure S4: Testing of PotB-ep18 and PotB-ep9 derivatives in minimal media. To determine whether the observed mutations altered ion selectivity we prepared swim plates in minimal agar in a range of phenamil concentrations (A: 0 μM phenamil, B: 20 μM phenamil, C: 100 μM phenamil) and with no sodium present (D/E: 85 mM KCl, 0 μM phenamil). Host: RP6665 (except E where indicated as RP437), 0.25% Difco Bactoagar, 10 mM $KPO_4$, 0.1% glycerol, 0.1 mM Thr, 0.1 mM Leu, 0.1 mM His, 0.1 mM Met, 0.1 mM Ser, 1 mM $MgSO_4$, 1 mM $(NH_2)_2SO_4$, 1 μg/mL Thiamine, 0.5% NaCl or KCl respectively (85 mM [NaCl]; [67 mM KCl]), 25 μg/mL chloramphenicol, phenamil as indicated. Incubated at 30°C for 24 hours.

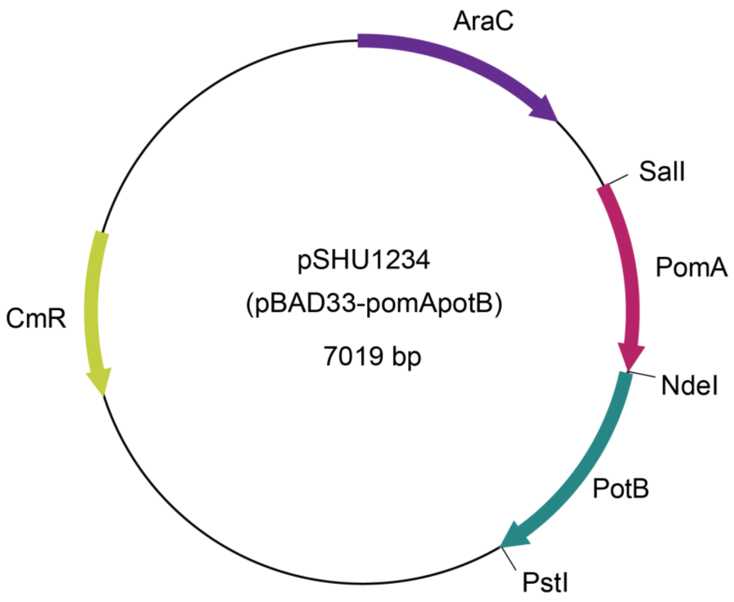

**Figure S5. Map of plasmid pSHU1234.** Plasmid based on pBAD33 backbone for expression of PomA and PotB with introduced NdeI site between PomA and PotB to aid in targeted random mutagenesis across PotB.

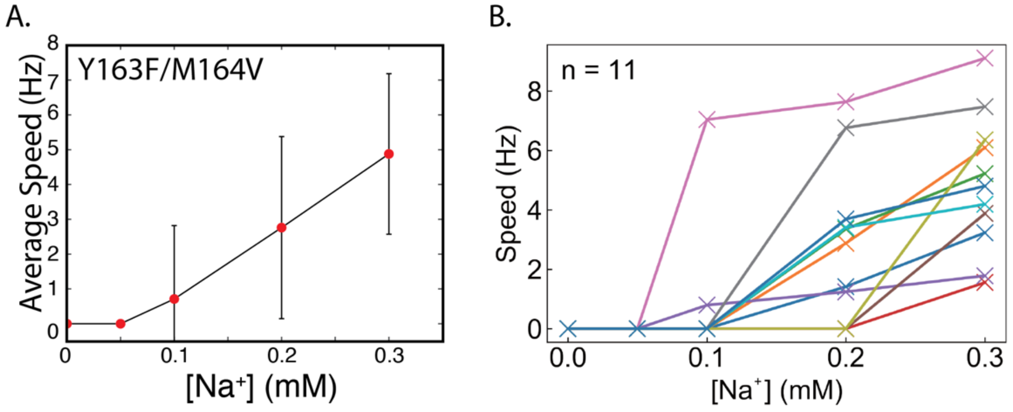

**Figure S6. Repeat measurements of Y163F/M164V at low sodium.** We repeated measurements of Y163F/M164V with finer spacing of sodium concentration and tracking single cell motility during sodium concentration transitions. (A) Mean and SD motility at 0, 0.05, 0.1, 0.2, and 0.3 mM $Na^+$. (B) Single cell motility tracked for each cell showing that all cells were motile at 0.3 mM $Na^+$, and 9 out of 11 cells were motile at 0.2 mM $Na^+$.

**Table S1**

| EP-PCR #ID | Mutant amino acids | swarm |
|---|---|---|
| PotB-ep1 | L15H, L35I, T60T, S73G, P74T, R154R, A207T, I265N, N269D | +++ |
| PotB-ep2 | I50T, M102V, D241G | + |
| PotB-ep3 | L15H, L17L, F33Y, M42K, R101S, S105G, K123N, L131P, E161E, N274S, T293T | ++ |
| PotB-ep4 | S38S, K100I, N180Y, L228S, H267R, E284V | ++ |
| PotB-ep5 | S105G, K132N, E268G, E284K | - |
| PotB-ep6 | S105G, L131P, K198K, P244L, H267L, S290G, V291V | + |
| PotB-ep7 | L15H, E41G, K109I, D113N, R128S, E138E, D166N, V230V, Q273P, S290G | ++ |
| PotB-ep8 | D9E, C10R, L36Q, V44I, M102L, I142I, I182I, E197D, I252F, E271D | +++ |
| PotB-ep9 | E88E, Y163F, M164V, L255M | ++ |
| PotB-ep10 | T21S, F39V, I68I, L124M, K198N, K226I, D245D, A286V | + |
| PotB-ep11 | G87G, S105S, L228F, P276P | + |
| PotB-ep12 | N94Y, R108W, R125R, L168L, F191L, L228F, P276T, Q288L, V291V | + |
| PotB-ep13 | F39L, F47L, Y163F, I171I, N176S, V248A, P283P, E284G, V289V | ++ |
| PotB-ep14 | M30L, F32I, T63S, G77G, I252F | +++ |
| PotB-ep15 | R128R, E262G, L266L, E284V | ++ |
| PotB-ep16 | D80E, I167V, P173H, E216V, L217M, Q288R, M294V | + |
| PotB-ep17 | V44A, K100I, K123R, N180D, I182I, E197G | + |
| PotB-ep18 | F22Y, L28Q, T56T, T60T, K100Q | +++ |
| PotB-ep19 | L36Q, G78S, I118T, E119V, L255L, Q260L, V289D, V291A | ++ |
| PotB-ep20 | V34A, T56T, T63A, D66D, E72E, E97K, K109N, D159V, N180S, A247A, K282K | ++ |

| | | |
|---|---|---|
| PotB-ep21 | L37H, E88D, L98L, Q137L, R165R, D190V, Q273P, P295Q | + |
| PotB-ep22 | P12P, L36Q, E88V, S105R, I118V, I145I, G196G, K198I, L217L, G220G, T236M, L239L, S240R, E281K, P283P | +++ |
| PotB-ep23 | Q92H, A172T, D223N, D245E, A286A | + |
| PotB-ep24 | N70Y, G112S, N180Y, G196G, R229R, Q260Q, I265I, L280M, V285V | ++ |
| PotB-ep25 | C8R, L28Q, M42V, F54Y, N70S, G196G, N202N, S224G, E268E, S290G, P295P | +++ |
| PotB-ep26 | C8Y, F22F, E96V, L107L, D113V, D159A, I171F, P173P, L217L, D223G, L228I, N258G, E262E, A286A, Q288L | + |
| PotB-ep27 | I50T, L117M, R125R, A234A, K259N, I265T | ++ |

**Table S1:** Mutation Tables for Phenamil-resistant outcomes from EP-PCR. All 27 selectable outcomes from EP-PCR are listed, with coding (red) and silent (black) mutations indicated in the second column. Comparative motility was recorded on a scale of 0-3 (-, +, ++, +++), with swim plate measurements shown in Fig. S1.

**Table S2**

| | Primer ID | 5>3 | Purpose |
|---|---|---|---|
| QuickChange | | | |
| | 1373 | TTATGGATGGGGACATACGCAGATTTGATGTC | PotB(F22Y)-F |
| | 1374 | GACATCAAATCTGCGTATGTCCCCATCCATAA | PotB(F22Y)-R |
| | 1375 | GCAGATTTGATGTCGCAGCTGATGTGTTTCTT | PotB(L28Q)-F |
| | 1376 | AAGAAACACATCAGCTGCGACATCAAATCTGC | PotB(L28Q)-R |
| | 1377 | ATCGAAGAGCTGAAACAACGCATGGAGCAAAG | PotB(K100Q)-F |
| | 1378 | CTTTGCTCCATGCGTTGTTTCAGCTCTTCGAT | PotB(K100Q)-R |
| | 1379 | GCCGATGTCGAACCCTTTATGCGCGACATTCT | PotB(Y163F)-F |
| | 1380 | AGAATGTCGCGCATAAAGGGTTCGACATCGGC | PotB(Y163F)-R |
| | 1381 | GATGTCGAACCCTATGTGCGCGACATTCTGCG | PotB(M164V)-F |
| | 1382 | CGCAGAATGTCGCGCACATAGGGTTCGACATC | PotB(M164V)-R |
| | 1383 | CGTCGCATCAGCCTGATGGTACTGAACAAACA | PotB(L255M)-F |
| | 1384 | TGTTTGTTCAGTACCATCAGGCTGATGCGACG | PotB(L255M)-R |
| | 1385 | GATGTCGAACCCTTTGTGCGCGACATTCTGCG | PotB(Y163F_M164V)-F |
| | 1386 | CGCAGAATGTCGCGCACAAAGGGTTCGACATC | PotB(Y163F_M164V)-R |
| Sequencing | | | |
| | 0363 | GCGTCACACTTTGCTATGCC | pBAD1212-f |
| | 0364 | TGGGACCACCGCGCTACTGC | pBADHind-r |
| | 0111 | GACATCGCGCTTACGGATGAAC | pomA382-403 |
| EP-PCR | | | |
| | 1176 | TAACTTGGAGAATTCATATGGATGATGAAGAT | Nde-potB-F |
| | 0104 | GAACTGCAGTCACCTCGGTTCGG | motAmotB-TR (Pst) |

**Table S2: List of primers used in this work.** Primers for QuickChange (point mutations to establish functional and non-functional mutations), Sequencing, and EP-PCR indicated respectively.

| Strain | Description | Reference |
|---|---|---|
| RP437 | Wild type for motility and chemotaxis:<br><br>F-, thr-1, araC14, leuB6(Am), fhuA31, lacY1, tsx-78, λ-, eda-50, hisG4(Oc), rfbC1, rpsL136(strR), xylA5, mtl-1, metF159(Am), thiE1 | (Parkinson and Houts, 1982) |
| RP6665 | Δ(*motA-motB*):<br><br>motA-motB)Δm12-13 his-4 metF(Am)159 (lac) Δu169 rpsL136<br>thi-1 ara-14 mtl-1 xyl-5 tonA31 tsx-78 | (Block et al., 1989) |

**Table S3: List of strains used in this work.**

**Non-phlyogenetic Logistic Regression (glm)**

| **PotB Position** | **r0_slope_Estimate** | **r0_slope_Std. Error** | **r0_slope_z value** | **r0_slope_Pr(>\|z\|)** |
|---|---|---|---|---|
| F22 | 2.947 | 1.300 | 2.266 | 0.023 |
| L28 | -18.817 | 3765.847 | -0.005 | 0.996 |

**Phlyogenetic Logistic Regression**
**(phyloglm with method = logistic_IG10)**

| **PotB Position** | **r2_slope_Estimate** | **r2_slope_StdErr** | **r2_slope_z.value** | **r2_slope_p.value** |
|---|---|---|---|---|
| F22 | 3.195 | 1.338 | 2.387 | 0.017 |
| L28 | 1.990 | 1.316 | 1.512 | 0.130 |

**Table S4: Correlation Analysis of Mutations at each respective site.** As per methods, we calculated a non-phylogenetic and phylogenetic general logistic model using packages glm and phyloglm in R respectively. Where the p-value of the slope is < 0.05, this indicates a significant correlation between the residue identity (ie F22 vs Y22) and PomB/MotB classification respectively.

| Strain | Ion Source |
|---|---|
| MotB_Escherichia_coli_strain_K12_MOTB_ECOLI | H+ |
| MotB_Salmonella_typhimurium_strain_LT2_MOTB_SALTY | H+ |
| MotB_Pseudomonas_aeruginosa_strain_ATCC_15692_Q9HUL2_PSEAE | H+ |
| MotB_Bacillus_subtilis_strain_168_MOTB_BACSU | H+ |
| MotB_Streptococcus_pneumoniae_A0A0T8PK69_STREE | H+ |
| MotB_Bacillus_licheniformis_strain_ATCC_14580_Q65KJ0_BACLD | H+ |
| MotB_Streptococcus_pneumoniae_A0A0E8TCW6_STREE | H+ |
| MotB_Helicobacter_pylori_strain_ATCC_700392_MOTB_HELPY | H+ |
| MotS_Oceanobacillus_iheyensis_A0A2P1WLE1_9BACI | Na+ |
| MotS_Bacillus_alcalophilus_G9I2I5_BACAO | Na+ |
| MotS_Bacillus_subtilis_subsp_natto_BEST195_BAI864791 | Na+ |
| MotS_Bacillus_licheniformis_A0A1Q9FXY5_BACLI | Na+ |
| MotD_Pseudomonas_aeruginosa_strain_ATCC_15692_G3XD90_PSEAE | H+ |
| MotB_Desulfovibrio_magneticus_strain_ATCC_700980_C4XPD2_DESMR | H+ |
| PomB_Vibrio_alginolyticus_O06874_VIBAL | Na+ |
| PomB_Vibrio_cholerae_serotype_O1_strain_ATCC_39315_Q9KTK9_VIBCH | Na+ |
| MotB_Aliivibrio_fischeri_KLU777421 | Na+ |
| PomB_Shewanella_oneidensis_MR_1_NP_7171461 | Na+ |
| MotB_Aquifex_aeolicus_strain_VF5_O67121_AQUAE | Na+ |

**Table S5: Classification of ion type used to calculate estimator.** After filtering (as per Methods) we were left with 19 strains where the ion type was identified from the literature. *Bacillus* (Fujinami et al., 2009); *E. coli* (Sowa and Berry, 2008)**,** *Salmonella* (Minamino and Imada, 2015)*; Pseudomonas* (Doyle et al., 2004)**;** *Aquifex, Shewanella* (Takekawa et al., 2015)**;** *Vibrio* (Atsumi et al., 1992); *Streptococcus* (Manson et al., 1977)**;** *Helicobacter* (Nakamura et al., 1998).

**Supplementary Note**

List of specific homologues from set of 948 Mot B homologues across *E. coli* K12's 308 residues that contained the mutations L28Q, Y163F, and M164V respectively. Both F22 and Y22 were present in many cases and as such is not listed.

**L28Q**
MotB *Myxococcus fulvus* A0A0F7BJE1 MYXFU

**Y163F**
MotB *Yersinia pestis biovar Orientalis* str IP275 A0A0M1V4N2 YERPE

**M164V**
*Betaproteobacteria bacterium* RBG 16 66 20 A0A1F3YFW8 9PROT
MotB *Caballeronia terrestris* A0A158KEN9 9BURK
*Paraburkholderia monticola* A0A149PEJ5 9BURK
MotB *Spirochaetae bacterium* HGW Spirochaetae 1 A0A2N1TQI3 9SPIR
*Spirochaetes bacterium* RBG 16 49 21 A0A1G3QXZ5 9SPIR
*Desulfuromonadales bacterium* C00003093 A0A1E7IF89 9DELT
*Oceanospirillum maris* UPI000403DD69
MotB *Oceanospirillales* A0A1H5Y5D3 9GAMM
TssL *Alteromonadaceae* bacterium A0A2D8J8H6 9ALTE
*Oceanospirillum maris* UPI000403DD69
MotB *Oceanospirillales* A0A1H5Y5D3 9GAMM
MotD *Gallionellales bacterium* GWA2 60 18 A0A1G0CPT2 9PROT
MotD unclassified *Nitrosomonadales* A0A1G0DLC6 9PROT
